# Patterns of microsatellite distribution reflect the evolution of biological complexity

**DOI:** 10.1101/253930

**Authors:** Surabhi Srivastava, Akshay Kumar Avvaru, Divya Tej Sowpati, Rakesh K Mishra

**Affiliations:** CSIR - Centre for Cellular and Molecular Biology, Uppal Road, Hyderabad 500007, India

**Keywords:** Microsatellites, SSR, density, length preference, evolution, phylogeny, taxonomy, simple sequence repeats, short tandem repeats

## Abstract

Microsatellites, also known as Simple Sequence Repeats (SSRs), are evolutionarily conserved repeat elements distributed non-randomly in all genomes. Many studies have investigated their pattern of occurrence in order to understand their role, but their identification has largely been non-exhaustive and limited to a few related species or model organisms. Here, we identify ~685 million microsatellites from 719 eukaryotes and analyze their evolutionary trends from protists to mammals. We document novel patterns uniquely demarcating closely related species, including in pathogens like *Leishmania* as well as in higher organisms such as *Drosophila*, birds, primates, and cereal crops. The distribution of SSRs in coding and non-coding regions reveals taxon-specific variations in their exonic, intronic and intergenic densities. We also show that specific SSRs accumulate at longer lengths in higher organisms indicating an evolutionary selection pressure. In general, we observe greater constraints in the SSR composition of multicellular organisms with complex cell types, while simpler organisms show more diversity. The conserved microsatellite trends and species-specific signatures identified in this study closely mirror phylogenetic relationships and we hypothesize that SSRs are integral components in speciation and the evolution of organismal complexity. The microsatellite dataset generated in this work provides a large number of candidates for functional analysis and unparalleled scope for understanding their roles across the evolutionary landscape.

## 1. Background

Repetitive DNA in eukaryotic genomes can be broadly classified into interspersed and tandem repeats. Microsatellites, also known as Simple Sequence Repeats or SSRs, are short tandem repeats of 1-6 nucleotide DNA motifs. They comprise a significant portion of the genome in complex organisms, often surpassing the proportion of coding sequences (Katti, et al. 2001). SSRs contribute to 3% of the human genome (Subramanian, et al. 2003), and display a non-random distribution in many genomes (Toth, et al. 2000; Katti, et al. 2001). They have high mutation rates due to polymerase slippage, with a bias towards elongation (Ellegren 2004). Due to their highly polymorphic nature, microsatellites have long been useful as molecular markers in a variety of fields including genotyping (Kashi, et al. 1997), marker-assisted selection (Cordeiro, et al. 2001), linkage analysis (Hearne, et al. 1992), and forensics (Zietkiewicz, et al. 1994). Though a majority of SSRs in genomes are present at intergenic and non-coding regions, a small proportion of SSRs occur within exons (Toth, et al. 2000; Li, et al. 2004). Abnormal expansion of SSRs within exons is associated with several diseases in humans such as Huntington’s disease and Spinocerebellar Ataxia (reviewed in (Usdin 2008)).

Recent studies have focused on the role of SSRs in important cellular functions such as epigenetic regulation of gene expression. AAG repeats were shown to be associated with repressive histone modifications (Greene, et al. 2007; Al-Mahdawi, et al. 2008), and H3K9me2 mediated regulation of the *Fxs* allele (leads to Fragile X Syndrome) is associated with expansion of CCG repeats at the FMR1 promoter in humans (Pietrobono, et al. 2005). SSRs present in proximal cis-regulatory regions such as promoters and introns have been shown to modulate gene expression via mechanisms that remain unclear (Bagshaw, et al. 2017). A few studies have also highlighted the roles of SSRs in genome organization. AGAT repeats of 40-48nt length are shown to function as enhancer-blockers in both *Drosophila* and human cells (Kumar, et al. 2013), while transcripts arising from AAGAG repeats have been identified as important constituents of nuclear matrix in *Drosophila* embryos (Pathak, et al. 2013).

Critical roles in modulation of gene expression and genome organization imply functional conservation across species. Indeed, microsatellites are believed to be under selection pressure, as apparent by their distribution and genomic abundance, which is much higher than expected by chance or random accumulation (Ellegren 2004). Despite being hypothesized to be the “tuning knobs” of evolution (Li, et al. 2004), a comprehensive analysis of these elements across the evolutionary landscape is lacking. A few studies (Toth, et al. 2000; Katti, et al. 2001) have chosen a small subset of representative species across evolution to analyze their SSR content. The caveat in such studies is that the results may reflect trends that are specific to the chosen species rather than the group they represent, particularly if the sequence quality of the available genomes is variable. Other studies (Hutter, et al. 1998; Morgante, et al. 2002; Ding, et al. 2017; Liu, et al. 2017) have limited their analysis to a single taxonomic group, making their observations difficult to understand in terms of the broader evolutionary landscape. Lastly, in silico SSR studies are limited by the efficiency, exhaustiveness and sensitivity of the various SSR identification programs they utilize and can be compromised by the quality of the SSR datasets generated (Lim, et al. 2013; Avvaru, Sowpati, et al. 2017).

Here, we have analyzed the evolutionary trends of microsatellite distribution across 15 taxonomic subgroups from protists to mammals. We have used a comprehensive SSR mining tool, PERF, to identify microsatellites in 719 eukaryotic species with 100% accuracy (Avvaru, Sowpati, et al. 2017) and discovered a large number of novel taxon-specific SSR enrichment patterns. We document evolutionary differences in SSR abundance in the context of genome size, GC content, and k-mer size of the motifs. Interestingly, microsatellite distribution trends accurately reflect phylogeny and we posit that they can be useful in understanding evolutionary relationships. Finally, we have used the available genome annotation data from 334 species to understand the distribution of SSRs in coding and non-coding regions. We discuss these findings in terms of the evolution of biological complexity, and report several remarkable observations that open new avenues for further experimental scrutiny.

## 2. Results

### 2.1. Overview of SSR distribution

We utilized our exhaustive repeat finding algorithm, PERF (Avvaru, Sowpati, et al. 2017) to search for all 501 possible SSR motifs occurring in 719 eukaryotic organisms for which genome sequence is available in the NCBI database (see Methods). We identified a total of 684,885,656 perfect SSRs (length >= 12bp) and analyzed their distribution patterns across evolutionary groups (Additional File 1). The organisms were divided into 5 main groups (protists, plants, fungi, invertebrates and vertebrates) constituting 15 subgroups. Figure 1 summarizes the distribution of SSRs across each taxonomic group and their genomic relationships. We find that simpler eukaryotes like protists and green algae (average of 0.1 million SSRs per genome) as well as fungi (average of 0.03 million SSRs per organism) have a much lower abundance of SSRs compared to plants (0.93 million SSRs per organism) and mammals (3.6 million SSRs per organism).

**Figure 1:**
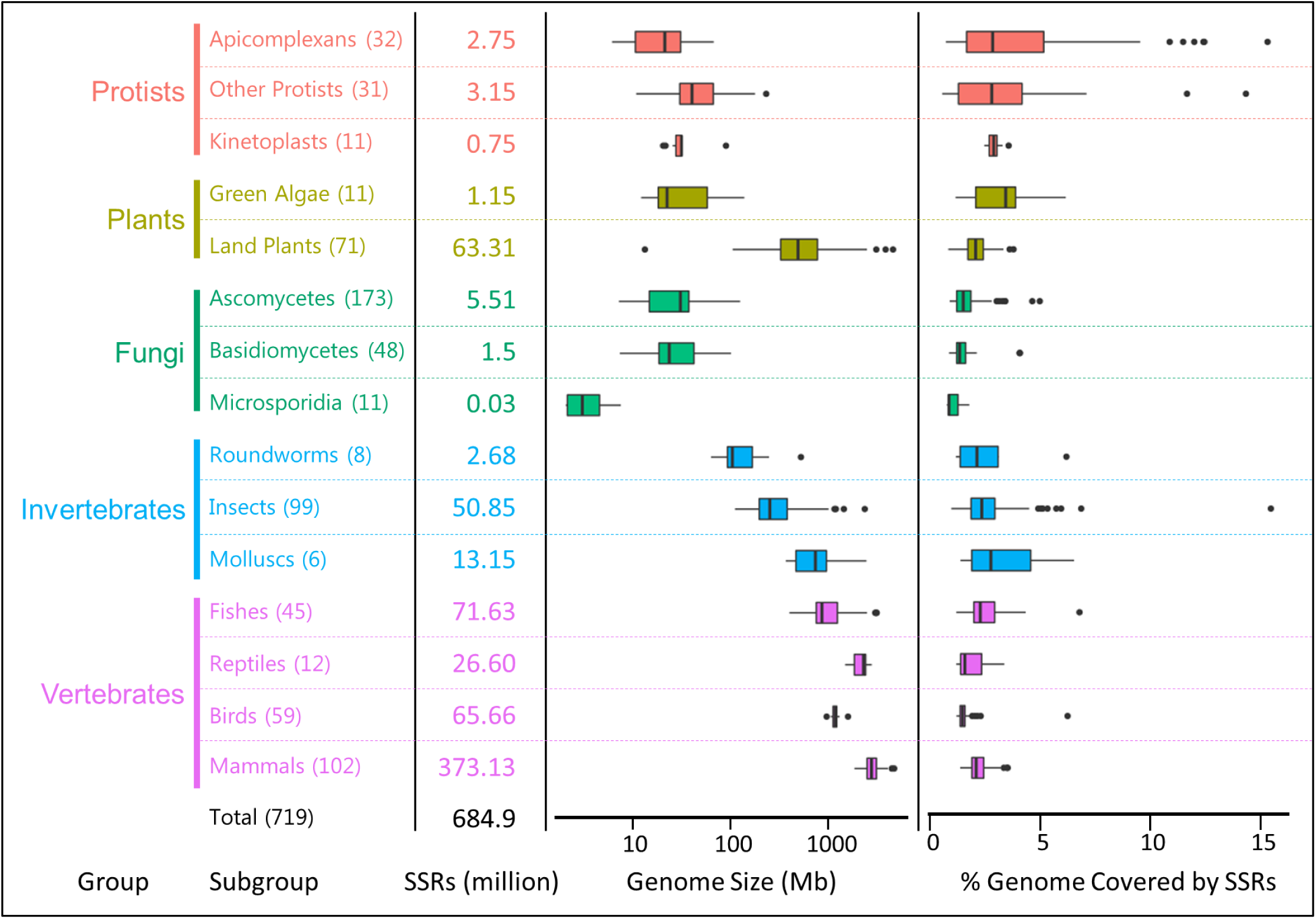
Overview of SSRs analyzed in this study. Approximately 685 million perfect SSRs (at least 12 bp in length) were identified from 719 eukaryotes across 15 subgroups, color coded and divided into5 groups. Numbers in parenthesis indicate the number of organisms in each subgroup. The total number of SSRs (in millions) analyzed in each subgroup is indicated. Box plots indicate the distribution of genomic sizes (highly variable across taxonomic groups, left) and the SSR coverage (% of genome covered by SSRs, relatively uniform, right) in each subgroup.

The total SSR abundance is correlated with genome size (Figure 2A, Pearson, r = 0.96). The top 50 organisms with high SSR frequency are mostly mammals including humans (4.6 million SSRs), a plant (*Aegilops tauschii*) and some fish (salmon species and coelacanth), all with 4-5 million SSRs and with genome sizes > 2.2 Gb. Of note, the highest number of SSRs are found in the octopus (7.6 million; genome size 2.3 Gb), the only non-vertebrate in the top 50 of the SSR abundance table (Additional File 1). At 1181 SSRs, the fungus *Encephalitozoon romaleae* (subgroup microsporidia) has the lowest number of SSRs, which correlates well with the fact that this pathogen has one of the smallest genomes studied (2.2 Mb) (Additional File 1). Since we wished to analyze patterns of SSR distribution over a wide range of genomic sizes, we felt that SSR abundance would not be a good measure to compare SSR trends across organisms in the evolutionary spectrum. Instead we looked at the density of SSRs (i.e. bp covered by SSRs per Mb of the genome) in order to normalize their occurrence to the genome size. We found that unlike SSR abundance, there is no correlation between the SSR density and the genome size (Figure 2B, Pearson, r = -0.04), though land plants do show a slight negative correlation (Pearson, r = -0.43) as documented previously (Morgante, et al. 2002).

**Figure 2:**
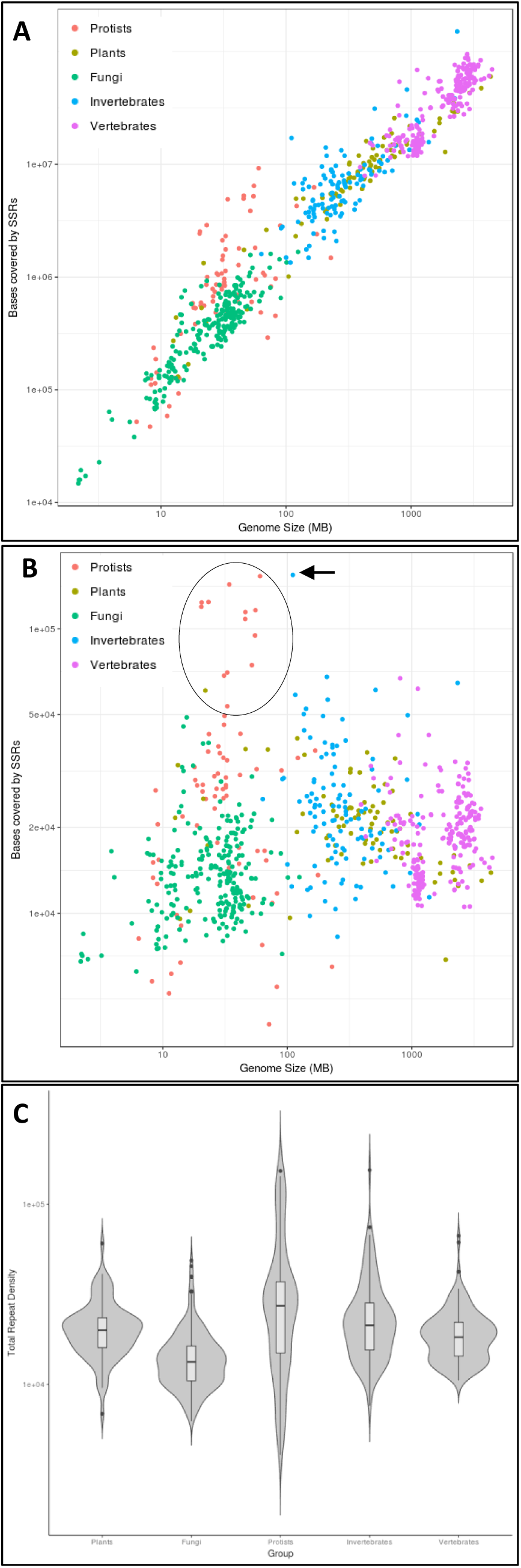
SSR attributes across the 5 groups. **A)** Abundance of SSRs (total bases covered by SSRs) correlates well with genome size (Pearson, r = 0.96), and therefore provides a biased view of SSR abundance in organisms with differing genome sizes. **B)** SSR density (bp covered by SSRs per Mb of the genome), on the other hand, shows no correlation with genome size (right; Pearson, r = -0.04) and is, therefore, a better measure to compare SSR distribution across organisms. Protists with very high density of SSRs are circled. The highest density of SSRs is found in the human parasitic louse (*Pediculus humanus corporis*, arrow). Colors indicate the 5 main groups of organisms analyzed. X axis: Genome size (Mb) in log scale; Y-axis: density of SSRs per MB of genome in log scale. **C)** Violin plots depicting the distribution of SSR densities across the 5 groups of organisms (X-axis). Protists show the highest variance compared to other groups (p <2.2e-16, F-test). The upper tail of the invertebrates group is because of a single outlier, the human parasitic louse (*Pediculus humanus corporis*), with an SSR density of154 Kb/Mb. Y-axis: density of SSRs per MB of genome in log scale.

We calculated the total SSR density (considering all the 501 repeat classes) for the 719 eukaryotes (Additional File 1). At a density of 154733 bp/Mb the human body louse (*Pediculus humanus corporis*) is the top ranked organism in terms of SSR abundance per Mb of the genome, i.e. 15.5% of its genome is covered by SSRs (Figure 2B, arrow). A recent analysis also made a similar observation, though from a comparison of only insect genomes (Ding, et al. 2017). At 21292 bp/Mb (SSR coverage 2.13%), humans have almost an order of magnitude lower SSR density than their parasitic louse. Among mammals, the kangaroo rat (*Dipodomys ordii*) has the highest density (34094 bp/Mb) followed closely by the house mouse (*Mus musculus*; 33776 bp/Mb). As a subgroup, mammals show little variance in SSR densities (about 3 fold difference between the highest and lowest SSR densities within mammals) (Table 1). Protists, on the other hand, occupy the highest density ranges and show the greatest variance among individual organismal SSR densities - upto 27 fold difference between the highest and lowest SSR densities among all protists (p < 2.2e-16, F Test, Figure 2C). For example, the *Dictyostelium* species (SSR density 143293 bp/Mb, SSR coverage 14.3%) and the *Plasmodium* species of the Apicomplexans subgroup (SSR density 124238 bp/Mb) have the highest SSR densities among all evolutionary groups examined while those belonging to the Oomycetes class and the *Entamoeba* species have significantly lower SSR densities such as *Giardia lamblia* (SSR density 5228 bp/Mb) and *Aphanomyces invadans* (SSR density 4070 bp/Mb, only 0.4% of the genome covered by SSRs); in fact they occupy the lowest end of the density spectrum among all 719 species (Additional File 1). The *Eimeria* species of protists from the Apicomplexans subgroup also have a high SSR density (average density 117719 bp/Mb) and these SSRs are notably GC-rich (SSR GC range of around 60%). It is interesting to note that these protists have such a high SSR density despite their small genome size (Figure 2B, circled). Fungi have the lowest average SSR density among all the subgroups while among higher vertebrates, birds have the lowest average, again with very little variance (Table 1), with the exception of the collared flycatcher (*Ficedula albicollis*) that has a very high SSR density of 61554 bp/Mb (SSR frequency of 1.27 million, 6.1% coverage in a genome size of 1.1 Gb).

**Table 1:**
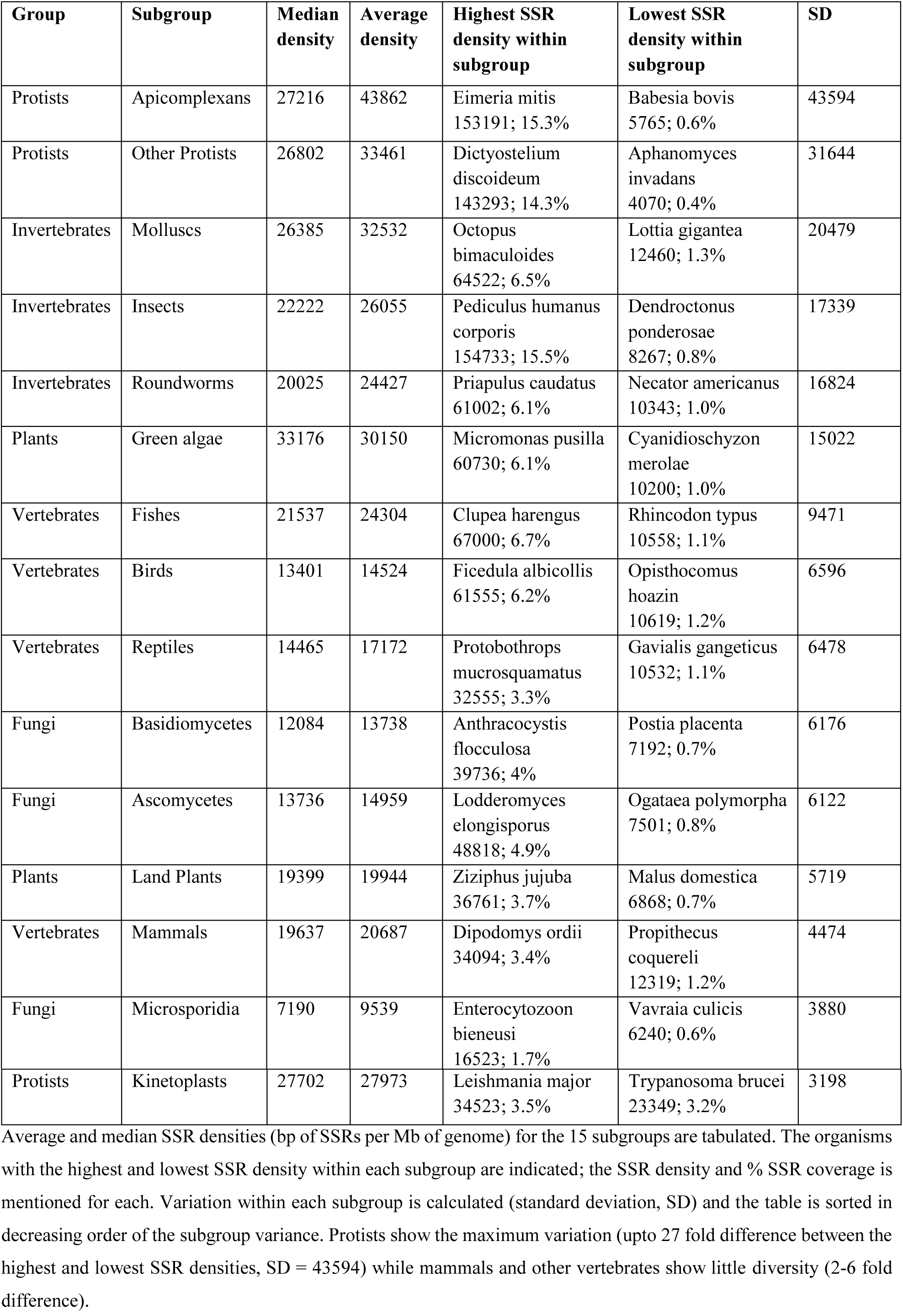
SSR densities across the 15 subgroups.

| Group | Subgroup | Median density | Average density | Highest SSR density within subgroup | Lowest SSR density within subgroup | SD |
| --- | --- | --- | --- | --- | --- | --- |
| Protists | Apicomplexans | 27216 | 43862 | Eimeria mitis<br>153191; 15.3% | Babesia bovis<br>5765; 0.6% | 43594 |
| Protists | Other Protists | 26802 | 33461 | Dictyostelium discoideum<br>143293; 14.3% | Aphanomyces invadans<br>4070; 0.4% | 31644 |
| Invertebrates | Molluscs | 26385 | 32532 | Octopus bimaculoides<br>64522; 6.5% | Lottia gigantea<br>12460; 1.3% | 20479 |
| Invertebrates | Insects | 22222 | 26055 | Pediculus humanus corporis<br>154733; 15.5% | Dendroctonus ponderosae<br>8267; 0.8% | 17339 |
| Invertebrates | Roundworms | 20025 | 24427 | Priapulius caudatus<br>61002; 6.1% | Necator americanus<br>10343; 1.0% | 16824 |
| Plants | Green algae | 33176 | 30150 | Micromonas pusilla<br>60730; 6.1% | Cyanidioschyzon merolae<br>10200; 1.0% | 15022 |
| Vertebrates | Fishes | 21537 | 24304 | Clupea harengus<br>67000; 6.7% | Rhincodon typus<br>10558; 1.1% | 9471 |
| Vertebrates | Birds | 13401 | 14524 | Ficedula albicollis<br>61555; 6.2% | Opisthocomus hoazin<br>10619; 1.2% | 6596 |
| Vertebrates | Reptiles | 14465 | 17172 | Protophthorops mucrosquamatus<br>32555; 3.3% | Gavialis gangeticus<br>10532; 1.1% | 6478 |
| Fungi | Basidiomycetes | 12084 | 13738 | Anthracoystis flocculosa<br>39736; 4% | Postia placenta<br>7192; 0.7% | 6176 |
| Fungi | Ascomycetes | 13736 | 14959 | Lodderomyces elongisporus<br>48818; 4.9% | Ogataea polymorpha<br>7501; 0.8% | 6122 |
| Plants | Land Plants | 19399 | 19944 | Ziziphus jujuba<br>36761; 3.7% | Malus domestica<br>6868; 0.7% | 5719 |
| Vertebrates | Mammals | 19637 | 20687 | Dipodomys ordii<br>34094; 3.4% | Propithecus coquereli<br>12319; 1.2% | 4474 |
| Fungi | Microsporidia | 7190 | 9539 | Enterocytozoon bienersi<br>16523; 1.7% | Vavraia culicis<br>6240; 0.6% | 3880 |
| Protists | Kinetoplasts | 27702 | 27973 | Leishmania major<br>34523; 3.5% | Trypanosoma brucei<br>23349; 3.2% | 3198 |

Average and median SSR densities (bp of SSRs per Mb of genome) for the 15 subgroups are tabulated. The organisms with the highest and lowest SSR density within each subgroup are indicated; the SSR density and % SSR coverage is mentioned for each. Variation within each subgroup is calculated (standard deviation, SD) and the table is sorted in decreasing order of the subgroup variance. Protists show the maximum variation (upto 27 fold difference between the highest and lowest SSR densities, SD = 43594) while mammals and other vertebrates show little diversity (2-6 fold difference).

The GC% of the SSRs is correlated with the genomic GC content (Additional File 2, Figure S1, Pearson, r = 0.94) and shows some interesting subgroup-specific patterns (Figure 3). Green algae (olive green arrow) and a few protists (*Aureococcus anophagefferens, Emiliania huxleyi, Thecamonas trahens*) with GC-rich genomes (genomic GC 55-65%) have an abundance of GC-rich SSRs (SSR GC range 75 – 100%). SSRs of intermediate GC content (SSR GC range 50%) are abundant in fungi. Higher organisms, however, have uniformly AT-rich SSRs (SSR GC range < 25%) with the exception of the bird *Ficedula albicollis* that has highly GC-rich SSRs (SSR GC content 77.8%, genomic GC 44.2%, Figure 3, pink arrow). We note that there is no correlation between the overall SSR abundance of an organism and its genomic GC content (Additional File 2, Figure S2).

**Figure 3:**
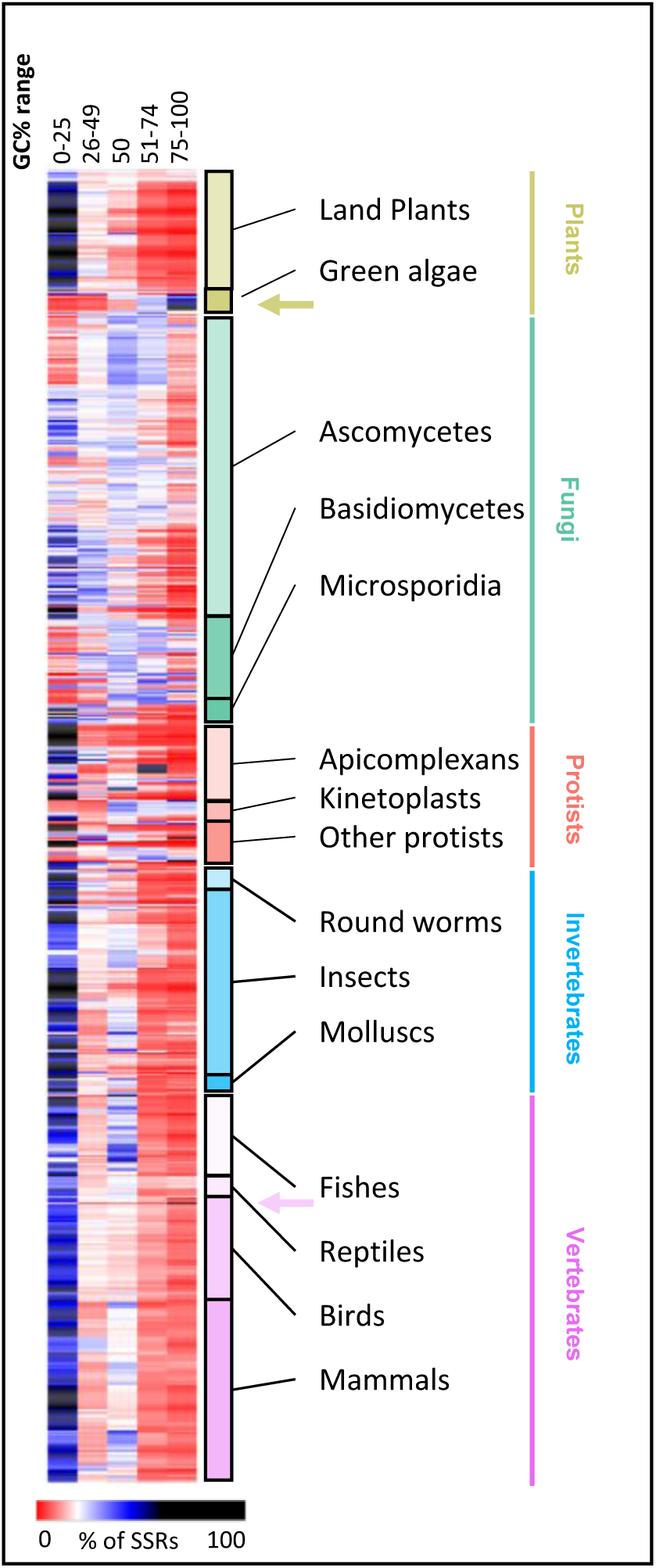
Subgroup-specific patterns of GC content in SSRs. SSRs are categorized by the GC percent of the repeat motif, calculated as defined below. Columns represent the GC% ranges and rows represent the 719 organisms analyzed. Color of the tile represents the percentage of SSRs falling within a given GC range (column) in that organism (row) as per the color scale shown below the heatmap. Subgroups are indicated on the right and grouped into the 5 main groups indicated on the right. Olive green arrow marks the position of green algae that are enriched for GC-rich SSRs and pink arrow marks the collared flycatcher *Ficedula albicollis* that has highly GC-rich SSRs (GC >75%). GC percent of SSR = (Number of G/Cresidues in the repeat motif *100) / length of the repeat motif.

### 2.2 SSR abundance trends across evolution

We plotted a ranked SSR density heat map (Figure 4, see methods) to look at density-based abundance trends of the 501 SSRs (columns) across all the 719 genomes (rows). SSRs were considered abundant in an organism if they occurred in the top 10 (black tiles) or top 25 (blue tiles) ranks. We discovered clear patterns of abundance that were distinct for different groups and even subgroups of organisms. As seen along the left-most columns of the heat map (Figure 4, black tiles at A1-K1 on the grid), a few SSRs are highly abundant across most organisms - viz C, AC, AG and the polyA repeat classes including A, A(n)T/G/C (density > 100 bp/Mb). But they are rare in green algae and some of the fungi of the ascomycetes and basidiomycetes groups; some of these SSRs are in fact entirely missing in these groups as indicated by the red squares (frequency < 10; B1 and B2 on the grid in Figure 4). Fungi have highly abundant ACG, CCG and other GC-rich repeats (black/blue tiles at B8-E8) that are not very abundant in higher organisms. Green algae show a high abundance of some GC-rich repeats (B8, B18, B20) correlating with their high average genomic GC content of 61.3%. Interestingly, in many fungi, especially from the Ascomycetes subgroup, upto 95% of the repeat classes appear to be missing (D2-D17, red tiles; frequency of occurrence < 10) though this trend shows a sharp change in the Basidiomycetes subgroup where only a few SSR classes are absent (E3 and E9-E12, red tiles). Some of the Protists (row F, red tiles) and green algae (B2-B7, B9-B17) too show a similar trend. Other than these species, most of the organisms (486/719) have some representation of all 501 classes of SSRs. Notably, the ACGCGT repeat is absent in about 64% of the organisms across evolution (A10.5 – G10.5 and H10.5 – K10.5, red arrow), except for bees, ants, wasps, and some fungi. Similarly, AGCGCT (red arrow) also appears to be unpopular, missing from about 51% of the genomes, including all vertebrates. Both these repeats in fact have the smallest lengths of occurrence as well (see below), suggesting that they are not well tolerated in genomes.

**Figure 4:**
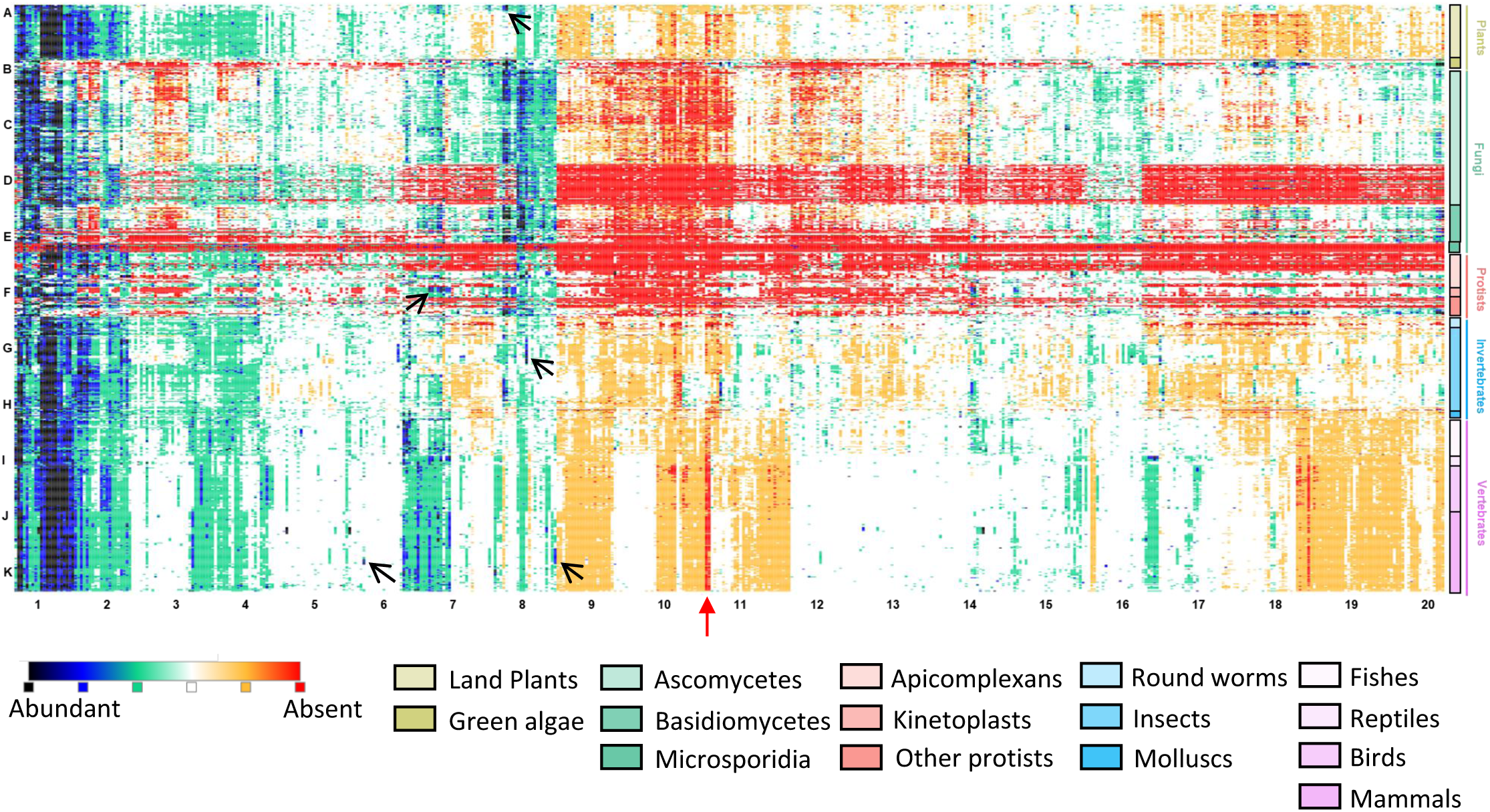
Enrichment trend of the 501 SSRs across 719 genomes. SSRs in an organism are ranked based on their density, defined as the total number of bp covered by the SSR in an organism normalized to genome size (that is, bp covered/Mb). The density based ranking is used to generate a heat map as per the color scale indicated. The 501 SSRs are arranged in columns and the 719 organisms are arranged in rows (the 5 main groups are indicated on the right; subgroups are indicated below the heatmap). Alphabets on the left and numbers at the bottom of the heatmap indicate approximate positions of the tiles in a virtual grid (eg., A1 to K20). Arrows mark the positions for uniquely abundant (enriched) SSR signatures described in the text and in table 2.

We next looked for SSRs that were highly abundant in only specific species or subgroups but not in any of the other organisms (Table 2). For example, a small group of land plants - cereals like maize, sorghum, millet, rice and corn (A8, arrow) - show some GC-rich repeats to be uniquely abundant. They also harbor abundant ACG and CCG repeats (A8) otherwise seen in fungi. Unique species-specific enrichment patterns can also be found in the *Leishmania* (F7) and *Drosophila* (G8, arrow) species as well as in higher organisms such as birds (I4), ruminants (bison, cattle, water buffalo, yak, sheep and goats - J5, J6, J14), and in primates (K6, AATGG and K9, ACCTCC, arrows).

**Table 2:**
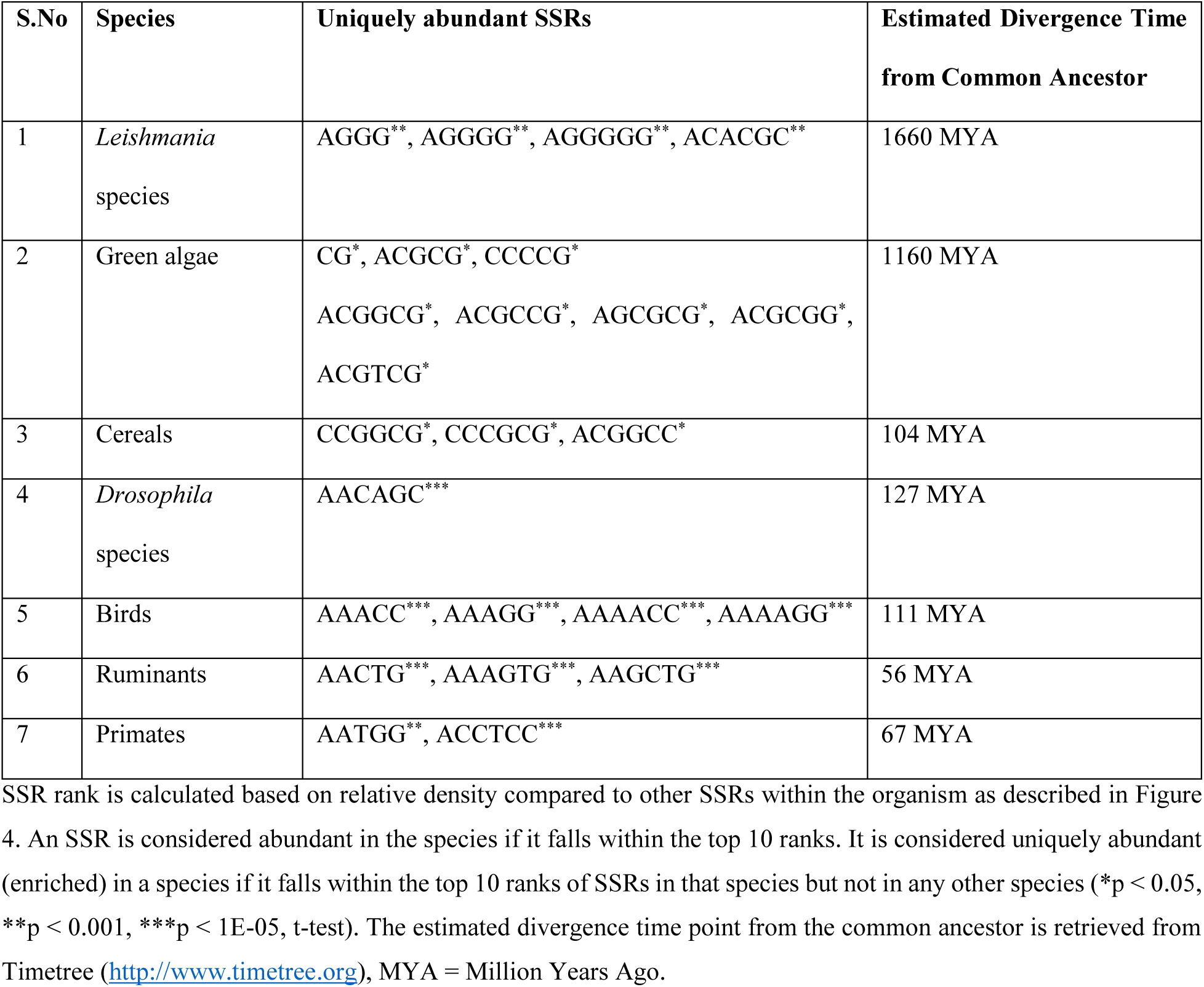
Uniquely abundant SSRs showing species-specific enrichment.

Further analysis of these signatures reveals interesting insights; for example, the ACCTCC repeat is in fact specific only to simians (median density 360 bp/Mb), enriched in all twenty simian species including humans but not in the four non-simian primates in our study (Additional File 2, Figure S3A). Interestingly, it occurs at a fixed length of 12 bp (Additional File 2, Figure S3B) and only in the sequence (ACCTCC)2 and not any other of its 5 cyclical variations (Additional File 2, Figure S3C). All other species, including close relatives of simians (Tarsiiformes and Strepsirrhini) have ACCTCC at a 10-fold lower median density of 36.3 bp/Mb, and all cyclical variations are represented in their genomes. These results implicate ACCTCC in a highly specific functional role in simians. We also notice other trends, for instance crocodilians such as gharials, crocodiles and alligators show a specific abundance for AGCCCC (I16) but many of these are not uniquely abundant or significantly enriched (p>0.05) and are hence not analyzed further. Notably, the abundance trend for specific SSRs is sharply contained within each of these groups of organisms (Figure 4), clearly defining them as species-specific signatures. A few of these SSRs have also been previously identified as being enriched in a species-specific manner in smaller scale studies limited to a few species or subgroups (Galindo, et al. 2009; Qi, et al. 2015; Qi, et al. 2016).

### 2.3 Length ranges of SSRs

We looked at the length of each SSR across all occurrences in the 719 organisms. As expected, longer SSRs were found to be present in the larger genomes (Figure 5, Spearman, r = 0.87). We identified the longest SSR found in each of the 719 organisms; AACCCT, the known telomeric repeat, is the longest SSR in 9% of the organisms (70 out of 743), with a top length of 15 kb in the fish *Rhincodon typus* (whale shark). We next looked at other longest repeats apart from the telomeric repeat and found that AT and AAT also frequently appear as the longest SSRs (in 8% organisms; top lengths 17.3 kb and 19.3 kb, respectively). The longest SSR seen among all organisms is almost 52 kb long – 12980 perfectly repeating units of AAAT in the mammalian *Cercocebus atys* (maps to an intergenic region, at a distance of 23.5 kb from the nearest gene, LOC105598351).

**Figure 5:**
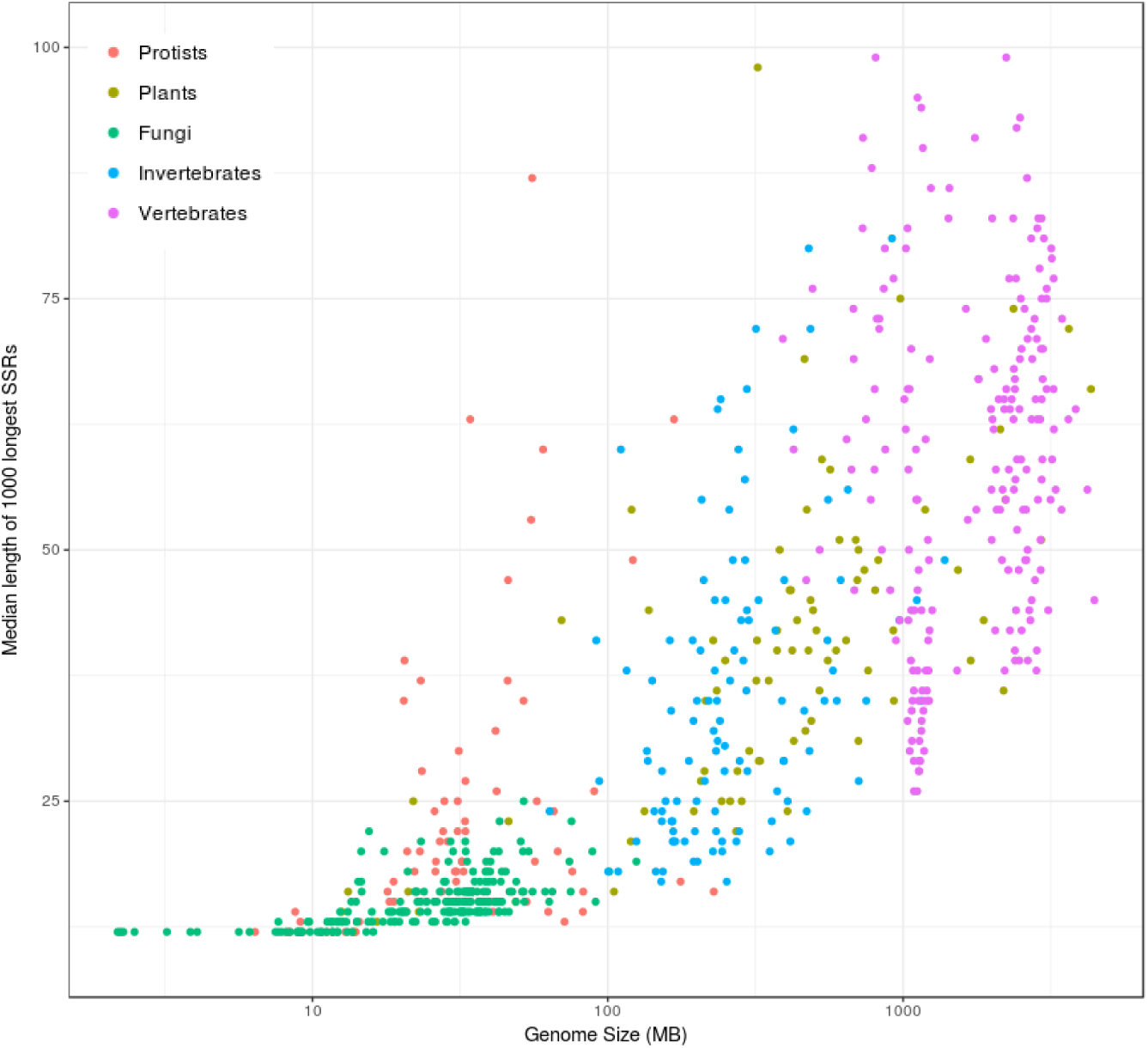
SSR lengths vs genome size. Scatterplot indicating the correlation between length of SSRs and the genome size of an organism (Spearman, r = 0.87). X-axis indicates the genome size in Mb on a log scale. Y-axis is the median length (bp) of the 1000 longest repeats present in an organism. The Y-axis is in log scale, and is trimmed to a maximum of100 to remove a single outlier – *Ficedula albicollis* (collared flycatcher), which has a median length of1897 bp. Colors of the dots indicate their group.

We then analyzed the length distribution for each SSR in a subgroup-specific manner (Figure 6A). As a single instance of a long repeat could potentially be an outlier, and may not accurately reflect the general trend, we derived the 100 longest repeat instances from each organism, and grouped such instances by the subgroup of the organism. Figure 6A depicts the range of subgroup-wise longest instances. Differences among subgroups within a group could often be correlated with the differences in their genome sizes. However, birds are an exception to this: despite having genome sizes significantly smaller than reptiles (p < 10e-9, pairwise t test, Bonferroni corrected) and mammals (p < 2e-16, pairwise t test, Bonferroni corrected), their SSR length range is higher than reptiles and mammals (p < 2e-16, pairwise t test, Bonferroni corrected).

**Figure 6:**
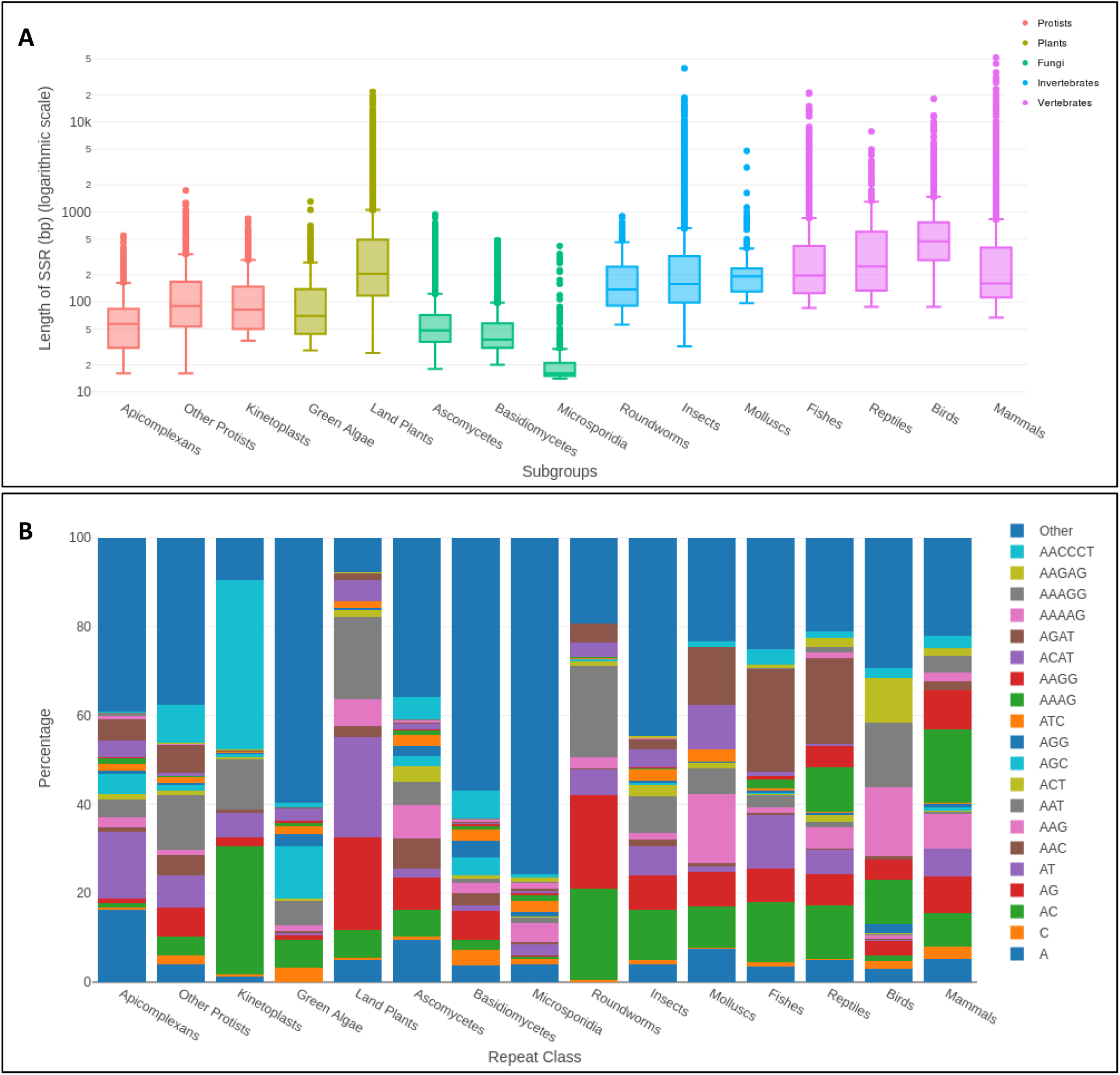
SSR lengths across subgroups. **A)** The top100 longest repeat instances from each organism are recorded separately and the data for organisms falling in the same subgroup are then grouped. Box plots showing the range of the longest100 repeat lengths in the organisms of a particular subgroup are plotted. Color indicates the main group to which the subgroup belongs. **B)** The percentage of occurrence of a repeat class in the top100 longest SSRs across all organisms is calculated to select SSRs that occurred at least1% of the times. The average percentage of these ‘highest occurring’ SSRs across organisms in each subgroup is plotted (Y-axis), categorizing the rest of the SSRs into “Other”. Individual SSRs are color-coded as indicated. The 15 subgroups are labeled along the X-axis.

We further checked if the longest instances of repeats in a subgroup belonged to specific repeat classes. We observed that distinct SSRs preferentially exist as long repeating units in different subgroups (Figure 6B). AACCCT, the telomeric repeat, is surprisingly not the longest repeat to be maximally represented in any subgroup except kinetoplasts (p < 10e-5, Fisher’s exact test). Instead, AGAT is the predominantly represented long SSR in fish and reptiles, and AAAG in mammals. Dimers of AC, AG and AT vary in their representation as long stretches in different subgroups; plants have maximal representation of AG and AT as longest SSRs while animals generally have longer AC stretches than AG or AT stretches (p < 10e-5, Fisher’s exact test). We next calculated the median from the 1000 longest instances of each SSR class to check for the preference of various SSR classes to exist as long repeats (Additional File 3). Interestingly, the median values range from as high as 4 kb (C repeats) to as low as 15 bp (AGCGCT repeats) (Table 3.1). While no obvious common motif pattern could be observed for the classes that showed the longest lengths, SSRs which showed the shortest lengths appeared to be of motifs with GC-rich centers flanked by A/T, but never completely GC-rich (Additional File 2, Figure S4), such as the two rarest repeats described in the previous section, AGCGCT and ACGCGT.

**Table 3.1:**
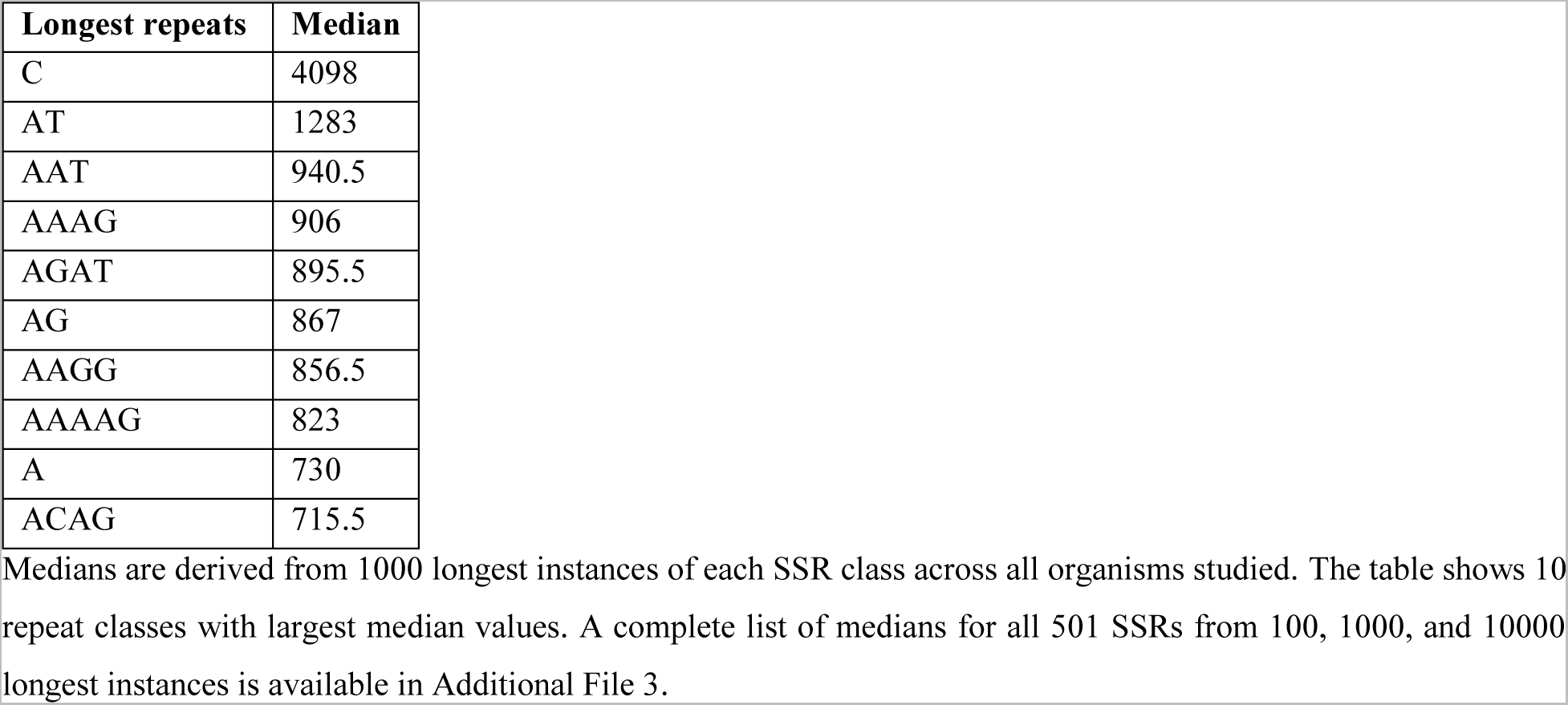
Median lengths of 10 longest SSR classes.

| Longest repeats | Median |
| --- | --- |
| C | 4098 |
| AT | 1283 |
| AAT | 940.5 |
| AAAG | 906 |
| AGAT | 895.5 |
| AG | 867 |
| AAGG | 856.5 |
| AAAAG | 823 |
| A | 730 |
| ACAG | 715.5 |

Medians are derived from 1000 longest instances of each SSR class across all organisms studied. The table shows 10 repeat classes with largest median values. A complete list of medians for all 501 SSRs from 100, 1000, and 10000 longest instances is available in Additional File 3.

The next question to emerge from this analysis was whether the longest repeats were present across all the organisms or if certain species preferentially accumulated instances of long SSRs. We therefore looked at the 1000 longest instances of each repeat class and checked if they belonged to the same species. Intriguingly, for many repeat classes, the longest instances did repeatedly occur in the same species. The most extreme case was again of the C repeats, where 974 out of the 1000 longest instances were from the collared flycatcher, *Ficedula albicollis* (Table 3.2). In fact, a few species have accumulated long instances of multiple repeat classes: the Asian tiger mosquito, *Aedes albopictus* was the most represented species for 28 different repeat classes, followed by the mouse, *Mus musculus* (26 classes).

**Table 3.2:** Preferential enrichment of long repeats in species.

| Repeat | Species | Number of instances out of 1000 longest |
| --- | --- | --- |
| C | *Ficedula_albicollis* | 974 |
| AAAATT | *Aedes_albopictus* | 953 |
| ACTCGG | *Selaginella_moellendorffii* | 923 |
| AATCCT | *Lupinus_angustifolius* | 864 |
| AGATCT | *Aplysia_californica* | 835 |
| AAAGC | *Aedes_albopictus* | 833 |
| AACGG | *Gorilla_gorilla_gorilla* | 801 |
| ACTAG | *Aedes_albopictus* | 799 |
| AAGATG | *Rhagoletis_zephyria* | 770 |
| AGGATG | *Lupinus_angustifolius* | 758 |

For every repeat class, the species to which each of the 1000 longest instances belonged was recorded. From this list, the number of times each species appears was derived, and the species that appeared the maximum number of times was designated as the most represented species. The top 10 repeat classes where more than 70% of the 1000 longest repeats belong to the same species are listed in this table. For the complete list of all 501 SSR classes, see Additional File 3.

The frequency of an SSR is expected to decline with increasing repeat length. We have demonstrated earlier that certain SSRs, however, show a preference for occurring at higher lengths in some organisms (Ramamoorthy, et al. 2014). A length preference is therefore defined as a sudden increase in the frequency of occurrence seen at a particular range of SSR length (Additional File 2, Figure S5). Our earlier work has indicated that 45 bp repeat size is the optimum length for a majority of the SSRs, especially in the human genome (Ramamoorthy, et al. 2014). Here we confirm that length preference is seen for relatively longer SSR size ranges in all genomes (~50 bp), except in fungi where SSR preferred lengths are generally smaller (~20 bp) (Table 3.3). The range of the preferred SSR lengths seen in each subgroup is tabulated in table 3.3. We find that only 131 out of the 501 SSRs can be associated with a specific length preference in any organism (see Figure 7 for a list of these 131 SSRs). We tabulated the length preference of each of the 131 SSRs across all 719 organisms and converted it to a heat map of percentage of organisms within a subgroup that show length preference for a given SSR (Figure 7). Fungi do not seem to have any preferred SSR lengths; none of the microsporidia show a length preference for any SSR (0%, red cells) while just 10-12% of Basidiomycetes show a preference for the polyA and polyC SSRs. Most lower organisms in fact are not associated with a length preference in any SSR though there is a significant increase in organisms showing SSR length preference in the higher subgroups (insects and beyond); mammals have a length preference associated with 85 of the 131 SSRs (last column, Figure 7). AAAG and AGAT show a length preference in >80% of mammalian species, followed by polyA and AAGG (73% mammals show length preference). The preference for specific SSR lengths observed here is not a function of genome size (Additional File 2, Figure S6).

**Figure 7:**
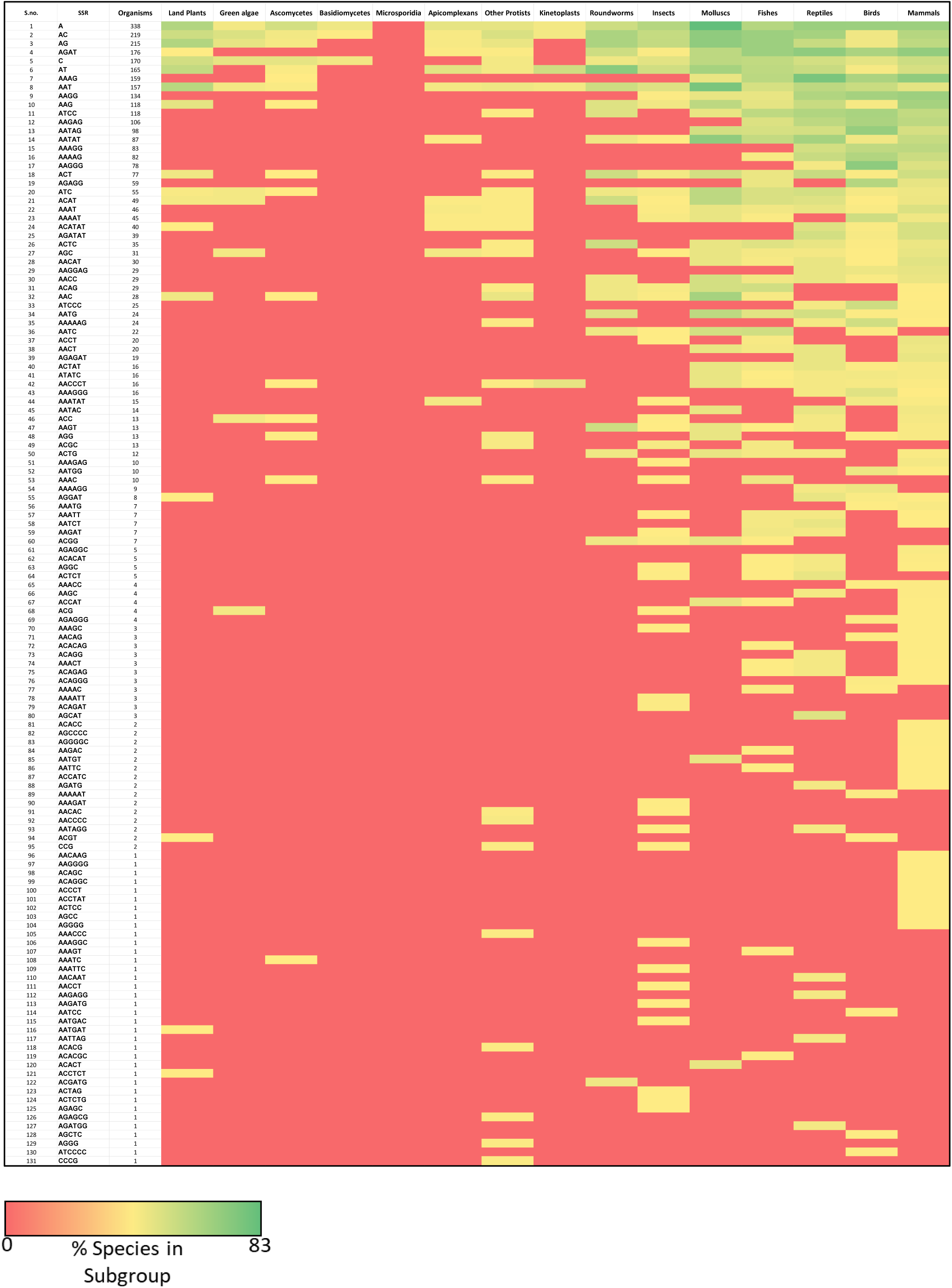
Summary of genomes that show SSR length preference. The 131 SSRs that show a length preference in any organism are arranged in rows. The number of organisms (totaled across all subgroups) that show a length preference for each SSR are indicated in the 3^rd^ column. The column headers indicate the 15 subgroups. The percentage of organisms in each subgroup that show a length preference for each SSR is indicated as a heatmap. Higher organisms prefer specific longer repeat lengths, for example many mammals show a length preference for a majority of the SSRs (for85 SSRs out of131).

**Table 3.3.** Length preference ranges shown by repeats in subgroups.

| **Group** | **Subgroup** | **Length preference peak start-end (bp)** |
| --- | --- | --- |
| Protists | Apicomplexans | 44 - 56 |
|  | Other Protists | 56 - 69 |
|  | Kinetoplasts | 35 - 46 |
| Plants | Green Algae | 48 - 64 |
|  | Land Plants | 46 - 58 |
| Fungi | Ascomycetes | 21 - 27 |
|  | Basidiomycetes | 16 - 21 |
|  | Microsporidia | NA |
| Invertebrates | Roundworms | 57 - 70 |
|  | Insects | 44 - 56 |
|  | Molluscs | 48 - 65 |
| Vertebrates | Fishes | 52 - 73 |
|  | Reptiles | 40 - 70 |
|  | Birds | 40 - 66 |
|  | Mammals | 36 - 60 |

Every instance of a repeat class showing preference for longer lengths in an organism was recorded (see Methods), and all organisms that belonged to the same subgroup were combined. The peak range of the subgroup was calculated by deriving the 5<sup>th</sup> percentile of peak starts and 95<sup>th</sup> percentile of peak ends within the subgroup to avoid outliers. NA indicates that not even a single instance of length preference was recorded for that subgroup.

### 2.4 Genomic patterns of SSRs by k-mer size

We also looked at the distribution of SSRs based on their k-mer size (1-6 bp) across all the subgroups (Figure 8). We find that monomers are abundant in birds and mammals (15-17% of total SSRs) and rare in green algae and fungi. Within mammals, primates have the highest contribution of monomers (18 %, Additional File 2, Figure S7A). Of the two possible monomers, polyA is largely preferred by all subgroups other than green algae (Additional File 2, Figure S7B), which is a reflection of their genomic GC content. The polyA bias is conspicuous especially in mammals, where A contributes >90% of the monomers (Additional File 2, Figure S7B); in primates, the proportion of polyA rises to 99% of all monomers (Additional File 2, Figure S8A). Birds and fungi have the lowest dimer content (Figure 8) while molluscs and fishes have the highest (median % = 20.93 and 20.52, respectively). Insects, mammals and land plants show similar abundance (~11%) but within mammals, rodents are prominent in their high dimer content (17.96% in rodents vs 11.8% in other mammals; Additional File 2, Figure S7A). CG repeats are extremely rare, contributing to less than 1% of all dimers in most species except green algae and basidiomycetes, where they contribute to 9.5% and 6% of dimers, respectively (Additional File 2, Figure S8B). AT repeats constitute the highest share of dimers in apicomplexans (93% of dimers) and land plants (62.3% of dimers), while AC dimers are the most frequent dimers in species of most other subgroups. Similar trends in dimer abundance have been previously documented for a few vertebrate species (Toth, et al. 2000).

**Figure 8:**
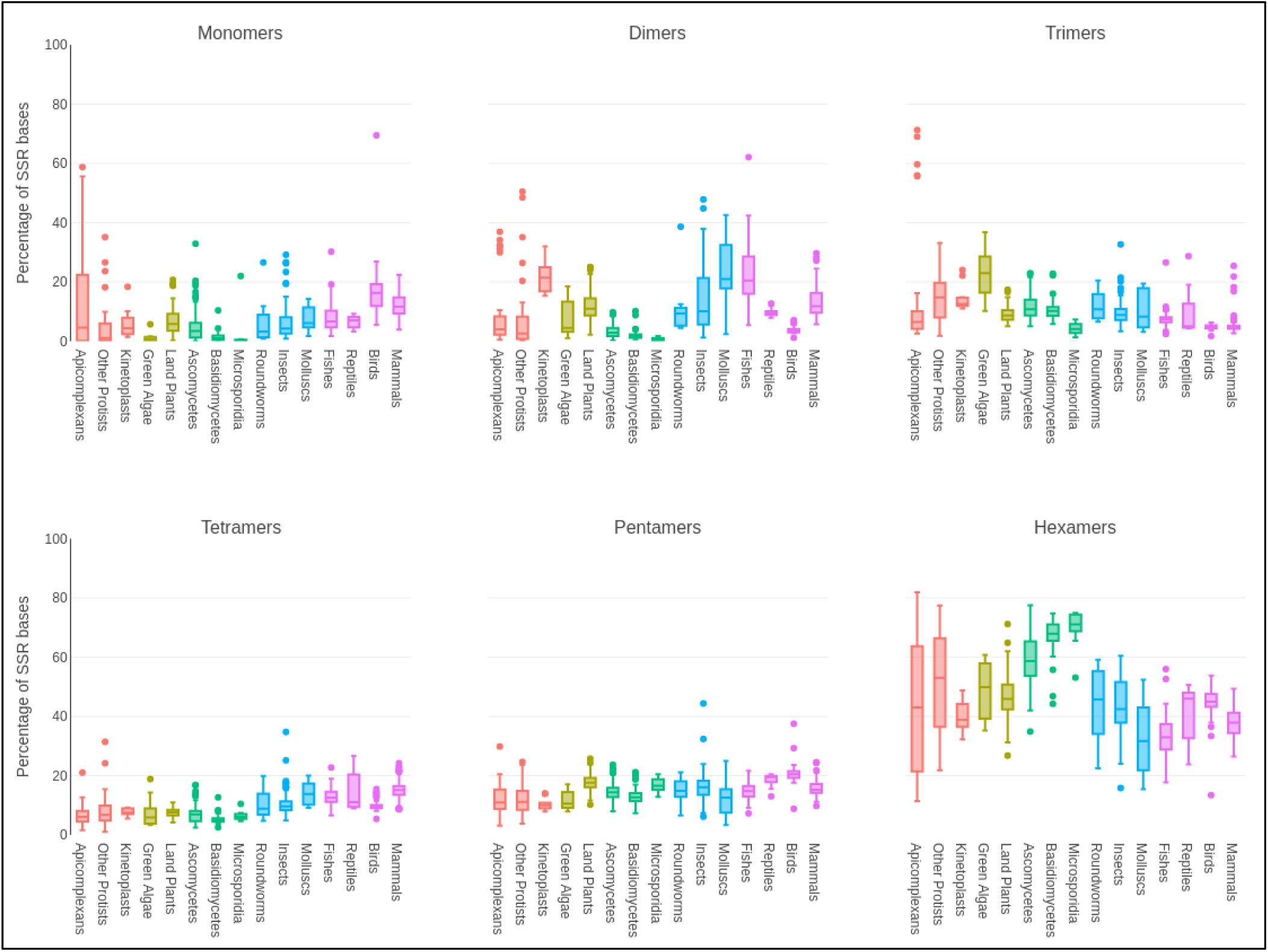
Composition of SSRs by their k-mer sizes. The number of bases covered by repeats in each k-mer class is summed across all organisms within a subgroup and divided by the total number of bases covered by all SSRs in that subgroup. Box plots for each subgroup represent the percentage of each k-mer base coverage (Y-axis) in the subgroup (X-axis).

Trimers are especially low in proportion in higher vertebrates such as birds and mammals but in green algae and some protists, trimers constitute a large proportion of all k-mer types, second only to their hexamer content (Figure 8). Overall, hexamers are the predominant SSR type in all organisms. Their proportion however is lower in higher organisms compared to protists, plants and fungi where they are the most abundant (~70% in microsporidia). Tetramers too show a noticeable difference, but in the opposite direction - tetramer percentages are lower in the lower eight subgroups (protists to fungi) compared to the last seven (invertebrates and vertebrates). Tetra‐ and pentamers show least variance in distribution across subgroups while dimers show the maximum divergence.

### 2.5 SSR distribution in genomic features

There has long been evidence for non-random genomic distribution of SSRs showing differential distribution across chromosomes and genomic features (Katti, et al. 2001). We, therefore, looked at the distribution of SSRs across 334 organisms for which genome annotations are available (Additional File 1) in order to understand biases in distribution trends and possible functions.

Overall, the distribution of SSRs in intergenic, intronic and exonic regions reflects their genomic distribution with intergenic regions having the highest abundance of SSRs (Figure 9A). However, we do see a small but significant underrepresentation of SSRs in exons (p < 10e-40, paired-sample t-test). This distribution remains the same across all the 7 subgroups, correlating with the respective genomic percentages of intergenic regions, introns and exons (Figure 9A). We checked for k-mer specific differences in the genomic distribution of SSRs. Introns and intergenic regions mostly show a similar distribution except in monomers which are slightly enriched in introns (Figure 9B). Exons, however, show a significant difference in the proportions of different k-mer types compared to the non-coding regions. Trimers and hexamers increase in exons compared to introns by ~7% and 16%, respectively (Figure 9B). An earlier study has indicated that trimers are the most abundant group in exons (Toth, et al. 2000), but we find that in fact hexamers tend to be more abundant in exons, similar to the trend observed in non-coding regions. Dimers and tetramers are under-represented in exons, occurring at higher frequencies in non-coding regions. This seems to imply that the underrepresentation of certain SSR k-mers is compensated by others in exons.

**Figure 9:**
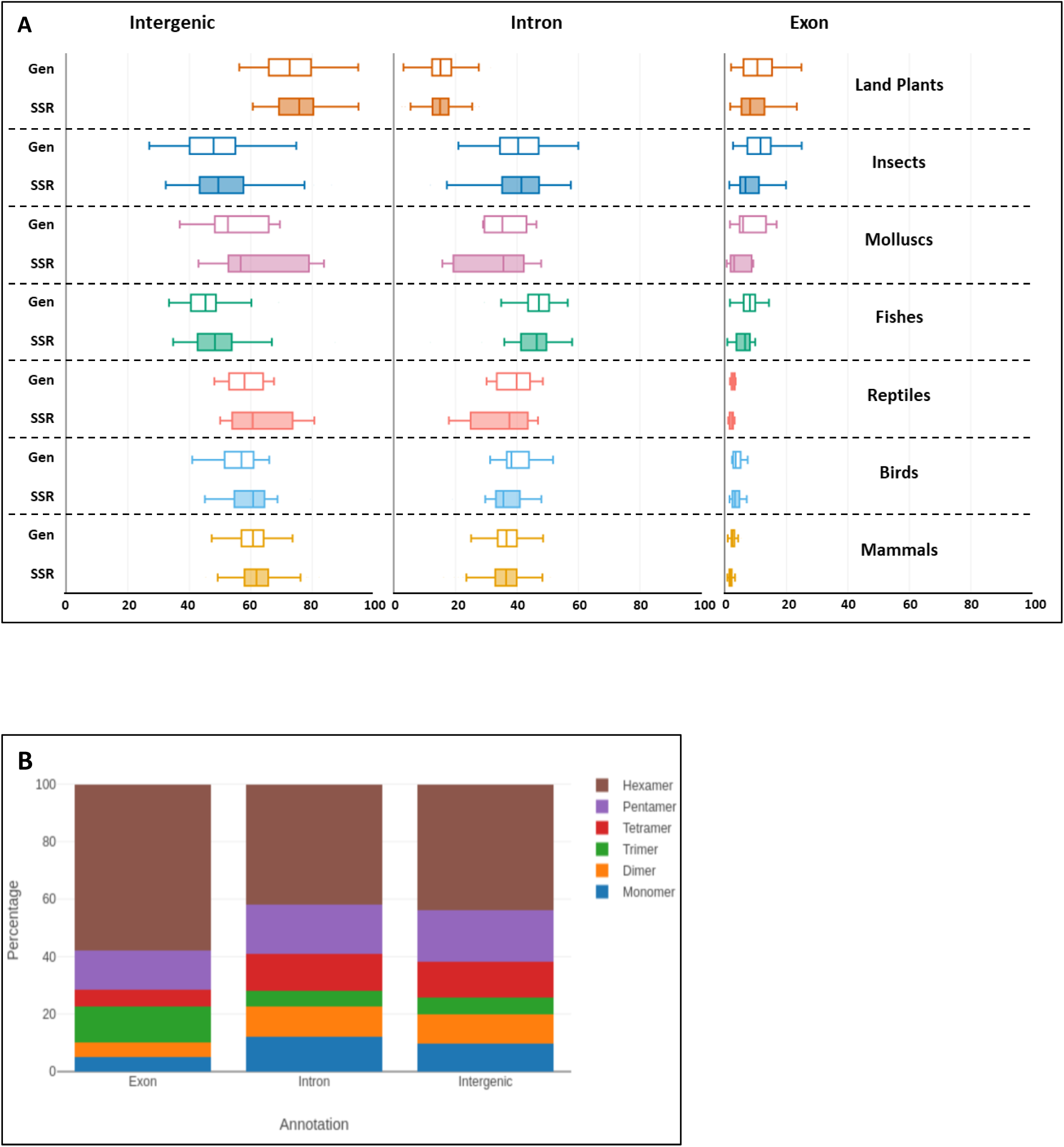
K-mer distribution of SSRs among various genomic features. **A)** Boxplots representing the proportion of SSRs in intergenic regions, introns, and exons across subgroups. The distribution of SSRs in intergenic and intronic regions (shaded boxes) mostly mirror their genomic distributions (non-shaded boxes). However, we see a small but significant underrepresentation of SSRs in exons (right panel, p <10e-40, paired-sample t-test). **B)** Average distribution of SSRs in coding and non-coding regions in all species studied, colored by k-mer size of repeat motif. The percentage of trimer and hexamer repeats is higher in exonic SSRs at the expense of tetramers and dimers. The fraction of each k-mer is calculated as the total bases covered by SSRs of a given k-mer size divided by the total bases covered by all SSRs. This value is multiplied by100 to derive percentages (the total for all k-mer sizes add up to100), represented on the Y-axis.

Subgroup specific k-mer distributions across genome annotations replicate these broad trends between exonic and non-coding regions (Additional File 2, Figure S9). Most repeat types such as monomers, dimers and tetramers are generally more abundant in introns and intergenic regions compared to exons across all taxa, while trimers and hexamers increase in exons. Mammals show an abundance of tetramers compared to trimers in introns and intergenic regions, a trend mirrored in other vertebrates albeit to a lesser extent. But the same is not seen in land plants and they have equivalent tetra‐ and trimers in their non-coding regions. Dimers are under-represented in exons across all groups, occurring at higher frequencies in non-coding regions. Interestingly, dimers are most enriched in the non-coding regions of fishes and molluscs contributing to 18% - 27% of SSRs, comparable to their hexamer content, while birds have the lowest contribution of dimers across all genomic regions (2-3%) as also mentioned earlier.

Lastly, we noticed that ACCTCC, the simian specific SSR identified in our analysis is under-represented in exons (p < 10e-11, paired sample t-test) and significantly enriched in intronic regions (p < 10e-15, paired sample t-test) (Additional File 2, Figure S10). This raises the intriguing possibility that it has a role in intronic splicing and/or acquisition of species specific splice variants. We found a similar trend in the genomic distribution of other uniquely abundant SSRs such as AACTG and AAAGTG in bovids, and AACAGC in *Drosophila* species. The proximity of such species-specific motifs to genes and *cis-*regulatory elements needs to be further analyzed to gain insights into their possible functions.

## 3. Discussion

Microsatellites are increasingly being recognized as critical sequence components with multiple roles in genome regulation. We identified perfect SSRs of 1-6 bp motifs from 719 genomes spanning 15 eukaryotic taxonomic subgroups including protists, plants, fungi, invertebrates and vertebrates. For 334 genomes for which annotation information is currently available, we also analyzed relative frequencies of these SSRs in genic and intergenic regions. We find that the distribution pattern of SSRs is a characteristic of the species or subgroup of the organism and that different taxonomic groups have distinct patterns of microsatellite presence and abundance. Our observations suggest the retention of specific SSR-based regulatory mechanisms as essential components of the genome.

### SSR constraints reflect evolutionary complexity

A greater degree of diversity in SSR patterns is observed in protists when compared to higher organisms (summarized in Figure 10). For instance, there are large variations in the densities of SSRs in protists, a subgroup that encompasses both SSR-dense as well as SSR-sparse genomes. Higher organisms, however, show lesser variation in SSR densities (Figure 10), suggesting greater constraints operating upon their genomes. Further, protists have very varied SSR GC contents as well - *Dictyostelium* and *Plasmodium* species harbor AT-rich SSRs while *Eimeria* and *Micromonas* species have GC-rich repeats. Green algae have a preponderance of GC-rich SSRs while most fungal SSRs are of intermediate GC content. On the other hand, higher organisms, including land plants, have a majority of genomes populated with only AT-rich SSRs with most GC-rich repeats having been filtered out (Figure 10). The stability in SSR composition in complex organisms is reflected in the relatively uniform content of genomic GC seen across vertebrates, while lower organisms show much more variable trends, reflecting a possibly dynamic usage, if any, of repeat sequences.

**Figure 10:**
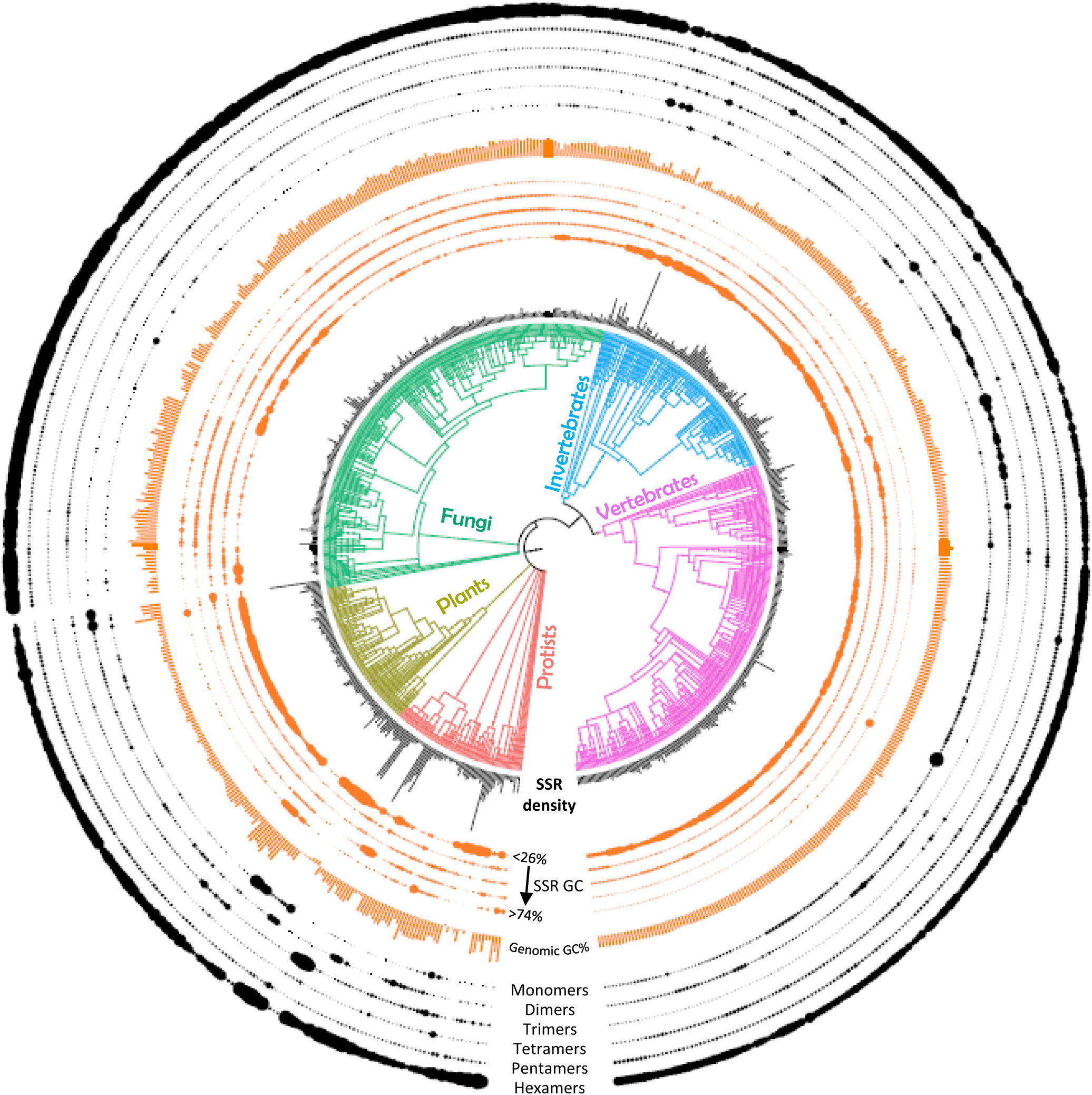
Phylogenetic tree representation summarizing attributes of all SSRs analyzed. The tree was constructed using iTOL (interactive Tree Of Life) webserver. The clade nodes are colored based on the 5 groups used in this study. Black bars (the innermost track) around the organisms represent the SSR density (bases covered per MB of genome) in each organism. The orange tracks around the SSR density show the SSR GC% in each organism (the innermost orange track represents the relative enrichment of motif s with <=25% GC, while the outermost orange track represents SSR GC >=75%) and the middle three tracks represent intermediate GC ranges. The size of each dot on the track (representing each organism) indicates the amount of SSRs present in that GC range. The orange bars represent the genomic GC content The black tracks show the k-mer distribution in each organism (the innermost black track represents monomers while the outermost black track represents hexamers). The size of each dot on the track (representing each organism) indicates the amount of SSRs present in that k-mer range.

### SSR enrichment shows species‐ and subgroup-specific trends

A unique aspect of this work is the analysis of SSR abundance trends from a phylogenetic perspective. Overall, SSRs are diverse in simpler organisms and their abundance appears to be relatively random compared to the enrichment trends shared by complex organisms. The sharp boundaries of changes in SSR abundance coincide perfectly with the evolutionary distinction between groups and subgroups. We were able to identify many taxonomic group specific microsatellite patterns within closely related species, for example the ACCTCC repeat specifically abundant in simians. Interestingly, we observe a very constrained selection for this motif (size not greater than 12 bp and no cyclical variations) which hints towards their possible role as regulatory motifs in simian genomes. Their significant enrichment in introns suggests that these elements could be playing a role in gene regulation via splicing and/or regulation of transcription rates.

AATGG is another primate specific signature identified in this work, albeit less significant when compared to ACCTCC. We have earlier shown AATGG to have remarkably different densities in human chromosomes, with an abundance in the Y chromosome (Subramanian, et al. 2003). An earlier microarray hybridization study has also shown a hominid (human, chimpanzee, gorilla, orangutan) specific enrichment for AATGG and suggested a role in gene regulation (Galindo, et al. 2009). Other specific patterns of enrichment also emerge from our analysis where a group of closely related species have clearly demarcated SSR densities. Such specific enrichment of SSRs could be causally linked to speciation. For instance, simians have a 10 fold greater abundance of the ACCTCC repeat compared to their close evolutionary cousins, the tarsiers and the strepsirrhines; simians diverged from the other primates around 67 MYA (million years ago). Similarly, the drosophilids, including *Drosophila melanogaster*, that share the uniquely abundant AACAGC signature repeat are 126 million years divergent from the other fruitflies of the family Tephritidae. It is possible that acquisition of specific repeats by a common ancestor millions of years ago offered a selective advantage to these species via novel SSR-mediated gene regulatory mechanisms. These need to be explored further in the context of protein factor binding, 3D long range interactions or direct transcriptional regulation of associated loci.

Interestingly, most fungi do not have >90% of the 501 possible SSRs, and also show the lowest SSR densities, indicating that these species may be under a different set of constraints in what their genomes can adopt. Other lower organisms – such as some protists and green algae – also lack 142
many repeat classes while invertebrates and vertebrates have a representation of almost all SSR classes. Exceptions are the ACGCGT and AGCGCT repeats that occur rarely in evolution, with >50% of all organisms excluding them altogether, including many vertebrates. Even in the few species of insects where these repeats occur at a moderate frequency, they are of short lengths. We hypothesize that such sequences constitute a rare class of SSRs that are intolerable in genomes.

### SSRs have subgroup-specific preferences for longer SSR lengths

Longer repeats are generally found in larger genomes and most repeats tend to decrease in frequency at higher lengths. PolyA-rich repeats such as AT and AAT and A(n)G tend to have the longest lengths of occurrence. Earlier studies (Toth, et al. 2000) have shown dimers to be the predominant longer repeats in introns and intergenic regions of genomes (except fungi). Subgroup specific longest SSRs identified in this work include AGAT (fish, reptiles) and AAAG repeats (mammals) as the predominant long repeats in vertebrates while AG/AC/AT dimers and AAT are frequent long repeats in lower organisms and in land plants (AT/AG). The differential enrichment for long lengths in SSR classes highlights subgroup-specific preferences where different regulatory mechanisms have been selected and fixed because they were advantageous.

We have observed that 131 out of 501 repeat classes show a specific enrichment at longer lengths in at least one species studied. The number of repeat classes that show this preference in an organism further correlates with their complexity: higher organisms show length preference in more repeat classes, maximally in mammals (65% out of 131 SSRs), while none of the fungi and only some protists show a length preference in any repeats. The preferred accumulation of longer repeats points to a selection pressure on these elements in a repeat class‐ and organism-specific manner. We have earlier shown that AGAT repeats show a preferred length of 40-48nt, and these elements function as enhancer blockers in *Drosophila* and human cells (Kumar, et al. 2013). Further, AGAT repeats of preferred lengths could bind to specific proteins more efficiently than repeats of smaller lengths. Hence, we speculate that repeats that show a length preference in various organisms might contribute to gene regulation via trans-acting protein partners. In this context, GATA repeat shows enrichment in many species (171 out of 719, out of which 82 are mammals), suggesting that its role could have been co-opted by multiple organisms.

However, this mode of regulation does not explain the possible functions of extremely long SSRs spanning kilobases, as protein binding generally does not span such lengths, even when cooperative binding is considered. Further, extremely long stretches (as recorded for the polyC repeat here) are likely to be fragile and disturb the nucleosome packaging, which is highly detrimental as seen in FragileX syndrome (Gatchel and Zoghbi 2005). Yet, we find that certain organisms have a capacity to accumulate and tolerate high repeat lengths. We have previously shown that AAGAG repeats get transcribed and are important constituents of nuclear matrix in *Drosophila* (Pathak, et al. 2013). Transcription from a few long non coding loci can trigger a large number of associated loci or cluster them into specific compartments, thereby acting as switches for coordinated genome regulation.

### SSRs operate under selection pressure

The fact that different SSRs seem to be preferred in a subgroup and species specific manner across evolution is a strong indicator of selection pressure and therefore the functional significance of microsatellites. Repeat elements have long been thought to have functional, mutational and evolutionary significance (McClintock 1950; Britten and Davidson 1969) and experimental evidence of a role for SSRs is increasingly available, albeit in only a few species per study (reviewed in (Kumar, et al. 2010)). The integrity of genome assembly and sequence information for different organisms can be an issue in such analysis, affecting SSR identification and inferences – in this context, our analysis of trends across subgroups alleviates, to an extent, the problems arising due to poor quality information from a particular genome. The consistent differences in SSR attributes between unicellular and multicellular organisms that we have described seem to point to a crucial role of SSRs in the creation and regulation of complex biological systems with multiple cell types. Perhaps different repeats need to be selected for functions in different cell types in multicellular organisms, and therefore constitute a basic mechanism for acquiring increasing complexity.

The observations presented in this work serve as a snapshot of evolution in the context of perfect SSRs – instances of repeats that were preserved without even a single nucleotide change. We have not considered imperfect SSRs in our study to avoid limitations posed by currently available imperfect SSR identification tools. Hence, this work cannot address whether these elements are a consequence of DNA repair / replication errors or mutational mechanisms. Many factors may be at play including, but not limited to, positive selection for repeat expansion (Li, et al. 2004), binding sites for proteins (Hu, et al. 2007; Liu, et al. 2017), effect on DNA structure and formation of secondary structures affecting transcription (reviewed in (Li, et al. 2002)), and relationships with high-order chromatin structure and genomic function (Kumar, et al. 2013; Pathak, et al. 2013). All these explanations notwithstanding, some repeats or repeat types and classes clearly show taxonomic group specific preferences in every attribute we have examined, representing their establishment in a common ancient founding member of all organisms in the group.

Our analysis reveals a few oddities worth mentioning. The genome of the collared flycatcher is unique in many aspects (Craig, et al. 2017). It is the only vertebrate to have such a high GC content of its SSRs despite an average genomic GC content of 44%. It is also unique in having the longest SSRs, with more than 15,000 instances of SSRs longer than 1 kb, out of which >13,000 are of C monomers. We also found abnormally high enrichment of select repeat classes in protists, viz. A repeats in *Plasmodium* species and AGC repeats in *Eimeria* species, which contribute to 40 - 70% of their total SSR content. An interesting observation is that most of these protists are parasites. Whether the preferential enrichment of a single repeat type in their genomes is beneficial to their pathogenesis remains to be explored further. In the light of the high polymorphism of microsatellites normally seen among individuals of the same species, identification of tightly conserved species-specific enrichment suggests a conserved functional role, especially prominent in higher organisms. Whether the specific enrichment indicates functional significance or mirrors close evolutionary relationships remains to be verified experimentally.

### The SSR paradigm of genome-wide regulatory mechanisms

As we move from simple (protists and fungi) to more complex (land plants, vertebrates and invertebrates) organisms, several aspects of SSR distribution become more selective, viz., density, GC content, length and k-mer composition. The evolution of complexity thus correlates with preferential and selective inclusion of SSRs that are retained for the evolutionary advantage they confer. The regulatory mechanisms involved may include protein factors that have evolved efficient binding to such repeats, perhaps in a cooperative manner or SSR transcription coordinating higher order regulation and long range interaction. Once acquired, these SSR mediated mechanisms can be easily scaled up to integrate a large number of loci (SSRs by definition being repetitive in nature) and play a role in the evolution of genetic programs that lead to the evolution of complexity. These are testable hypotheses and the SSR signatures identified in this work can serve as a starting point for understanding this paradigm.

## 4. Materials and methods

### Collection of genome data and evolutionary ordering of species

The latest versions of eukaryotic genome sequences available on NCBI’s RefSeq database were downloaded from the FTP site of NCBI (ftp://ftp.ncbi.nlm.nih.gov/genome). The hierarchical classification of each organism into kingdom, group and subgroup was adopted from NCBI’s “Genome Browse” table. Further taxonomic classification of organisms was gathered using an R package, *taxize*. We removed subgroups with less than 5 organisms from our analysis, after which we were left with 719 species spanning 15 subgroups (Figure 1). All the organisms and subgroups were arranged in an evolutionary order using *Time Tree* (http://www.timetree.org/), a web application for information on the evolutionary timescale of life. This order of organisms was consistently followed for all further analyses.

### Identification of SSRs

Perfect SSRs >= 12nt in length were identified from sequences of all downloaded genomes using a Python-based exhaustive algorithm, PERF (Avvaru, Sowpati, et al. 2017). The 5356 possible permutations of 1–6nt long DNA motifs were grouped into 501 unique classes of SSRs based on the cyclical variations and strand of the motif sequence, as described previously (Avvaru, Saxena, et al. 2017). A repeat class motif represents all the motifs which are cyclical variations of itself and of its reverse complement (Additional File 4). PERF reports all SSR locations in the genome in BED format, with additional columns describing the length of the repeat sequence, the repeat class, number of times the motif is repeated in tandem (repeating units), and the actual repeat motif (defined by the start of the SSR sequence, irrespective of repeat class). Using these parameters, we identified a total of 684,885,656 repeats from the genome data of 719 species.

### Calculation of basic SSR attributes

For each organism, we calculated a few parameters that outline the prevalence of SSRs in the genome. SSR frequency is the total number of SSRs found in the genome. The total bases covered by SSRs in the genome is calculated by summing the lengths of all the SSRs. To normalize for differences in genome sizes across evolutionary groups, we derived the SSR density for each genome, defined as the number of bases covered by SSRs per MB of genome. This was calculated by dividing the total SSR bases with the genome size in MB. We have used SSR density throughout the study, unless otherwise mentioned. The SSR GC% of an organism is the GC% of the sequence formed by concatenating all the repeat sequences found in the genome. A master table containing all the SSR attributes, along with the taxonomical classification and genome information of each organism, is provided as Additional File 1.

### Repeat class specific abundance trends across evolution

SSR frequency, base coverage, and density for each of the 501 repeat classes were calculated in each organism using in-house Python scripts. To identify repeats that are specifically enriched in various taxa, we ranked all the repeat classes based on their density in each organism. The ranks were further encoded with scores on a scale of 3 to -2 to improvise clustering. Briefly, we first gave the lowest score of -2 to those repeats which had a frequency of <10 in a given organism, to reduce sampling bias. Further, we assigned scores 3, 2, and 1 to repeats with the top 10, 25 and 100 ranks respectively. We have confirmed that the minimum density of SSRs in the top 100 ranks across all organisms is 12.04. Repeats in the bottom 100 ranks and frequency of at least 10 were given a score of -1. All other repeats were assigned a score of 0. A matrix was built using the score information, where each row represents an organism and columns represent the repeat classes. Hierarchical clustering of the repeat classes was done using the Euclidean distance between columns of the matrix. The clustered matrix was visualized as a heatmap using Morpheus (https://software.broadinstitute.org/morpheus/), an interactive tool for generation and exploratory analysis of heatmaps. The color scale on the heatmap ranged from a high score of 3 (black) to a lowest of -2 (red) as described above and in Figure 4.

### Length preference analysis

Contrary to the expected gradual decrease in abundance of longer repeats, some repeat classes show an increase in abundance at longer lengths. This pattern appears as a peak with a local maxima when unit length vs abundance for a repeat class is visualized as a line chart (Additional File 2, Figure S5). A custom Python script was developed to detect repeat classes showing this pattern in all organisms. The script compares the abundance at consecutive unit lengths to detect an increase. The length before the first detected increase is considered the peak start. The script further checks if the increase continues to a local maxima followed by a decrease in abundance. The endpoint of the curve is where the abundance goes lower than the abundance at the peak start. To filter false positives, we consider instances which span over at least 4 consecutive unit lengths (start and end included), and where the abundance at the initial increase unit is greater than 10.

### K-mer and GC composition of SSRs

Repeat classes are categorized based on the length of the repeat motif from monomers to hexamers. K-mer analysis involves calculation of different SSR attributes as described above for each k-mer category. The base coverage of a k-mer category is calculated by summing up the base coverage for all the repeat classes falling in that category. For GC composition analysis, we categorized the 501 repeat classes into 5 groups based on the GC content of their repeat motif. This was calculated using the 12bp string (minimum length cutoff) formed by repeating the base motif in tandem. The 5 groups of SSR GC content are <=25%, 26-49%, 50%, 51-74%, >=75%, which encompassed 70, 120, 133, 108, and 170 motifs respectively.

### Genomic annotation of SSRs

GFF files containing gene annotation information of various organisms were downloaded from the FTP site of NCBI (ftp://ftp.ncbi.nlm.nih.gov/genome). Annotation for SSRs was done based on the GFF files using an in-house Python script. In brief, the script uses the genic and exon coordinates to identify SSRs that overlap either exons, introns, or intergenic regions. For each SSR, the output includes its genomic annotation, whether it is promoter-associated, and its distance to the nearest TSS. In addition, for all exonic SSRs, the percentage overlap of SSR with an exon is reported. This was done to ensure that our results are not skewed because of a high proportion of SSRs falling within exon-intron boundaries. We verified that >95% of exonic SSRs show a complete overlap with exons.

### Statistical analysis

Two sample t-test was done using t.test() function in R. Pairwise calculations were done using pairwise.t.test() in R, and p-values were adjusted using Bonferroni correction. Paired sample t-test was done using ttest_rel() function of SciPy package in Python. Variation in SSR densities was measured using Standard Deviation of SSR densities in a subgroup, and F-test to assay the significance in variance was done using var.test() function in R. One-way ANOVA followed by Tukey’s post-hoc tests were done using aov() and TukeyHSD() functions of R respectively, using a confidence interval of 0.99. Fisher’s exact test was done using fisher.test() function of R. Most plots were made using ggplot2 and the Plotly API of R and Python.

## Additional Files

Additional file 1 (.xls, 109 KB): AdditionalFile1_mastersheet - Mastersheet of SSR attributes and genomic information for 719 organisms arranged in evolutionary order.

Additional file 2 (.pdf, 946 KB): AdditionalFile2_SupplFigs - Supplementary figures S1 – S10 along with their legends.

Additional file 3 (.xls, 98 KB): AdditionalFile3_length - Table of SSR‐ and species-wise median lengths of longest repeat instances.

Additional file 4 (.xls, 9 KB): AdditionalFile4_motifs - Table explaining the classification of SSR motifs. Repeat classification is shown for a normal motif (AAG), a palindrome (ACGT) and a cyclical variation of a palindrome (AGCTCG, cyclical variation of CTCGAG).

## Acknowledgements

We acknowledge Phanindhar Kundurthi for suggestions and critical reading of the manuscript. This work was supported by grants from the Council for Scientific and Industrial Research (BSC0121, BSC0118).

